# Whole-exome sequencing of plasma cell-free DNA portrays the somatic mutation landscape of refractory metastatic colorectal cancer and enables the discovery of mutated *KDR*/VEGFR2 receptors as modulators of anti-angiogenic therapies

**DOI:** 10.1101/177287

**Authors:** Rodrigo A. Toledo, Elena Garralda, Maria Mitsi, Tirso Pons, Jorge Monsech, Estela Vega, Álvaro Otero, Maria I. Albarran, Natalia Baños, Yolanda Durán, Victoria Bonilla, Francesca Sarno, Marta Camacho-Artacho, Tania Sanchez-Perez, Sofia Perea, Rafael Álvarez, Alba De Martino, Daniel Lietha, Carmen Blanco-Aparicio, Antonio Cubillo, Orlando Domínguez, Jorge L. Martínez-Torrecuadrada, Manuel Hidalgo

## Abstract

The non-invasive detection of cancer mutations is a breakthrough in oncology. Here, we applied whole-exome sequencing of matched germline and basal plasma cell-free DNA samples (WES-cfDNA) on a *RAS/BRAF/PIK3CA* wild-type metastatic colorectal cancer patient with primary resistance to standard treatment regimens including VEGFR inhibitors. Using WES-cfDNA, we could detect 73% (54/74) of the somatic mutations uncovered by WES-tumor including a variety of mutation types: frameshift (indels), missense, noncoding (splicing), and nonsense mutations. Additionally, WES-cfDNA discovered 14 high-confidence somatic mutations not identified by WES-tumor. Importantly, in the absence of the tumor specimen, WES-cfDNA could identify 68 of the 88 (77.3%) total mutations that could be identified by both techniques. Of tumor biology relevance, we identified the novel *KDR*/VEGFR2 L840F somatic mutation, which we showed was a clonal mutation event in this tumor. Comprehensive *in vitro* and *in vivo* functional assays confirmed that L840F causes strong resistance to anti-angiogenic drugs, whereas the *KDR*/VEGFR2 hot-spot mutant R1032Q confers sensitivity to cabozantinib. Moreover, we found a 1-3% of recurrent *KDR* somatic mutations across large and non-overlapping cancer sequencing projects, and the majority of these mutations were located in protein residues frequently mutated in other cancer-relevant kinases, such as EGFR, ABL1, and ALK, suggesting a functional role.

In summary, the current study highlights the capability of exomic sequencing of cfDNA from plasma of cancer patients as a powerful platform for somatic landscape analysis and discovery of resistance-associated cancer mutations. Because of its advantage to generate results highly concordant to those of tumor sequencing without the hurdle of conventional tumor biopsies, we anticipate that WES-cfDNA will become frequently used in oncology. Moreover, our study identified for the first-time *KDR*/VEGFR2 somatic mutations as potential genetic biomarkers of response to anti-angiogenic cancer therapies and will serve as reference for further studies on the topic.

## INTRODUCTION

Judah Folkman demonstrated almost half a century ago that neovascularization is a key requirement for solid tumor growth and metastasis^1^. This discovery motivated the development of anti-angiogenic drugs targeting cancer endothelial cells in order to deprive tumors from their blood supply^2^. Three decades later, bevacizumab, a recombinant humanized monoclonal antibody blocking vascular endothelial growth factor A (VEGF-A) and consequently, the activation of the VEGF receptor 2 (VEGFR2) on endothelial cells, became the first anti-angiogenic drug to be approved by the Food and Drug Administration (FDA) for cancer treatment^3^. The safety and efficacy of bevacizumab have been assessed in several randomized controlled clinical trials that showed an extended overall survival of patients with metastatic colorectal cancer (mCRC)^3,4^. However, the majority of patients (approximately 50% and 80% in first and second lines, respectively) do not benefit from such treatment; biomarkers predictive of the response to bevacizumab, as well as new agents targeting the VEGFR2 pathway, such as regorafenib, are still a fundamental unmet medical necessity^5^.

The possible involvement of genetic variants of the VEGF-VEGFR1/2 pathway in the outcome of anti-angiogenic treatment has been extensively investigated. Although some studies have suggested the potential association of tumor response with VEGF/VEGFRs germline polymorphisms^6^, these results could not be confirmed in subsequent assessments^7^. Recent *in vitro* studies demonstrated that VEGFR2 plays a functional role not only in endothelial cells, as is usually assumed, but also prominently in cancer cells^8^. This finding indicates that, similar to other drug target kinases such as the epidermal growth factor receptor (EGFR) and the ABL proto-oncogene 1 (ABL1)^9-11^, VEGF/VEGFR2 mutations occurring in the tumor cells, rather than inherited polymorphisms, could be regulating drug efficiency.

The conventional approach used for the discovery of the genetic causes of therapy resistance relies on genetic or genomic analyses of the patient’s tumor, which implies surgery or tumor biopsy. Here, we identified a novel somatic mutation in the key player of angiogenesis *KDR*/VEGFR2 (L840F) in an mCRC patient who was highly refractory to standard and experimental therapies, including angiogenesis inhibitors, despite wild-type (WT) results in targeted liquid biopsy analysis, by following an innovative approach of whole-exome sequencing of matched germline and plasma cfDNA (WES-cfDNA)^12^. *In vitro* and *in vivo* functional analyses demonstrated that this mutation, as well as other recurrent *KDR*/VEGFR2 mutations reported in large cancer databases, can modulate the efficiency of anti-angiogenic therapies.

## RESULTS

### CLINICAL

#### Case report

A 56-year old man was diagnosed in our center with a WT *KRAS*/*NRAS*/*PIK3CA*/*BRAF* mCRC cancer with liver and lung metastases (cT4N2M1). He received FOLFIRI-cetuximab as frontline treatment but there was evidence of tumor progression after the first tumor evaluation as suggested by the appearance of a new liver lesion. Consequently, he received the following treatments sequentially: FOLFOX-bevacizumab, afatinib with cetuximab, oncolytic adenovirus monotherapy (the last two lines of treatment in the context of phase 1 clinical trials), rechallenge with capecitabine-bevacizumab, and finally, regorafenib; however, there was no response and progressive clinical worsening occurred (Figures 1A and S1). The lack of radiological or clinical benefits in response to any of these treatments was mainly because of the persistent growth of his liver tumor burden, as the single lung metastasis grew less prominently. The patient died within a short time period (14 months) after the initial diagnosis due to complications of his progressive disease. Highly-sensitive BEAMing (beads, emulsions, amplification and magnetics; Sysmex Inostics, USA) digital PCR liquid biopsy assays revealed no alterations in the hot-spots of the *KRAS*/*NRAS*/*PIK3CA*/*BRAF* genes in the patient’s prior-and on-treatment plasma cell-free DNA (cfDNA) samples. Moreover, none of these mutations were identified in the tumor samples by Cobas (Roche), suggesting that other genes could be responsible for such prominent resistance. Therefore, we sought to design an unbiased strategy to identity and characterize newly implicated genes in cancer therapy resistance.

**Figure 1:**
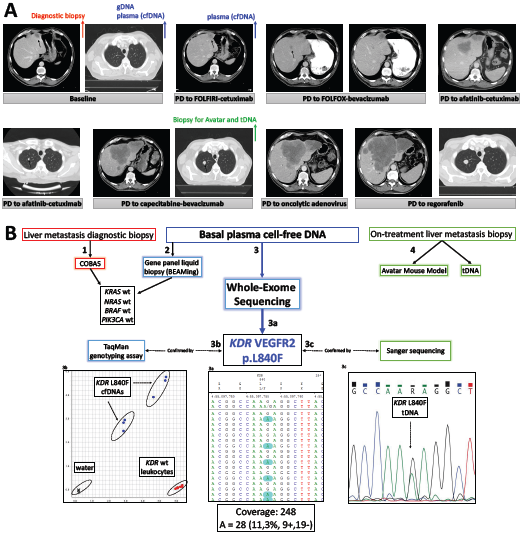
Discovery of the *KDR* /VEGFR2 L840F somatic mutation in a refractory mCRC patient. A) Thoracic computed tomography scans of the refractory mCRC patient. Imaging examinations show no response to any treatment and disease progression to FOLFIRI-cetuximab, FOLFOX-bevacizumab, afatinib-cetuximab, capecitabine-bevacizumab, oncolytic adenovirus and regorafenib. The time points of sample collection are pointed with colored arrows: red for diagnostic biopsy, blue for DNA sequencing, and green for Avatar generation and tumor DNA sequencing. Metastatic foci are indicated by black arrows. B) Non-invasive strategy for the discovery of primary therapy-resistance gene(s). Genetic and genomic analyses were performed according to the DNA sample availability. First, tumor biopsy was used (and exhausted) on routine genetic analysis that revealed no mutations in the main resistance-associated genes *KRAS*/*NRAS*/*BRAF*/*PIK3CA*. Second, whole-exome sequencing was performed using basal plasma cfDNA (to obtain both germline and cancer genetic variants/mutations) and matched leukocyte DNA (to filter out germline variants), and identified the *KDR*/VEGFR2 L840F somatic mutation. Third, the mutation was confirmed by Sanger and TaqMan assays in pre-and post-plasma cfDNA samples and in a second biopsy. A fragment of the biopsy was used to generated the patient-derived xenograft (PDX) Avatar model and for tumor DNA exome sequencing.

#### Samples

The first sample available for molecular analyses was a fragment from the patient’s liver metastasis obtained during the diagnostic biopsy, which was used (and exhausted) on the histopathology and the routine genetic analysis, owing to its small size. Following the diagnosis of *KRAS*/*NRAS*/*PIK3CA*/*BRAF* WT mCRC, the patient accepted to participate in this exploratory study. Peripheral whole-blood and basal plasma samples were collected and used to test the WES-cfDNA strategy. After progression to first line, a second blood/plasma sample was collected, and another liver metastasis biopsy was performed (Figures 1 and S1). The on-treatment plasma sample was used for validating the *KDR* mutation identified on basal plasma. The second liver biopsy was divided in two parts: one fraction was used to generate the patient-derived xenograft (PDX) model, herein called Avatar model, and the other fraction for tumor WES (WES-tDNA). The list of somatic mutations identified by WES-cfDNA was compared to that from WES-tumor to assess the mutation detection efficiency of WES-cfDNA.

### GENOMIC ANALYSES

#### Discovery of the clonal, somatic *KDR*/*VEGFR2* L840F mutation by WES-cfDNA

WES-cfDNA confirmed the WT status of the *KRAS/NRAS*/*PIK3CA*/*BRAF* genes (Figures S2-S4), as previously indicated by BEAMing. In addition, WES-cfDNA uncovered two known colorectal cancer driver mutations, *APC* c.3964G>T E1322X (COSM18702) and *TP53* c.659A>C Y220S (COSM43850) (Figures S5-S6), as well as the *KDR* c.2518C>T mutation leading to the L840F change in the VEGFR2 receptor (Figures 1B, S7). This *KDR* mutation was predicted to be pathogenic based on the high degree of conservation of L840, and had not been previously reported in cancer or population sequencing projects. We confirmed the *KDR* c.2518C>T allele in the patient´s basal and on-treatment cfDNA samples (collected after progression with FOLFIRI-cetuximab) by TaqMan genotyping assay, but not in the corresponding gDNA, confirming its somatic status (Figures 1B, S7-S8).

To estimate the clonality of *KDR*/VEGFR2 L840F, we used mutated allele frequencies (MAFs) data from WES of both tumor and plasma cfDNA (shown in Figure 2A and Table S1). Importantly, the MAFs of *KDR*/VEGFR2 L840F were similar to the MAFs of trunk CRC mutations, such as those in *APC* and *TP53*. The MAF of *KDR*/VEGFR2 L840F in the tumor sample was approximately 30%, whereas the MAFs for the *APC* p.E1322EX and *TP53* p.Y220SY mutations were both approximately 50%. Importantly, the MAF of *KDR*/VEGFR2 mutation in plasma (11%) was higher than the MAF of the *APC* mutation (8%), whereas the MAF of the *TP53* mutation was the highest (18%). Our data demonstrate that the *KDR*/VEGFR2 mutations can occur before and independently of targeted therapy pressure. Moreover, after normalization with the MAFs of *APC* and *TP53*, our results show that L840F was likely a clonal mutation event in this tumor (see (supplementary discussion, point I).

**Figure 2:**
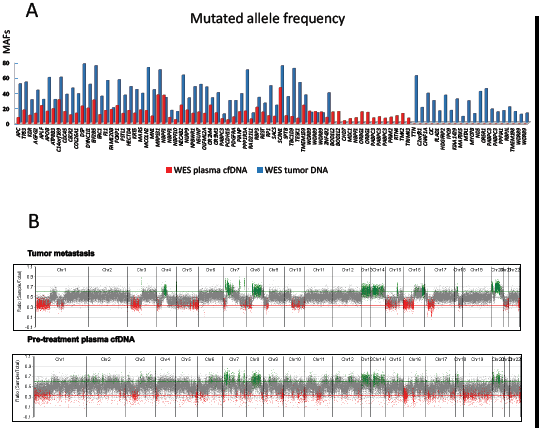
Concordance between the genomic landscape identified by whole-exome sequencing of plasma cfDNA and tumor DNA. A) The histograms represent all the genes with somatic mutations identified by plasma cfDNA whole-exome sequencing (in red) and by tumor whole-exome sequencing (in blue). The list and genomic annotation of all mutations are shown in Table S1. The mutated allele frequencies (MAFs) of each mutation in relation to all reads are depicted in Y. B) Copy number alteration landscape portrayed by the whole-exome sequencing of tumor DNA (upper part) and pre-treatment plasma cfDNA (lower part). Gains are depicted in green, losses in red, and normal (balanced) in grey.

#### High concordance between WES-cfDNA and WES-tumor

Once we gained access to the tDNA after the second liver metastasis, we confirmed the L840F mutation by Sanger sequencing (Figure 1B) and performed WES-tumor for comparison with WES-cfDNA. Applying the same stringent bioinformatics filters (described on the online methods), WES-cfDNA could detect 73% (54/74) of the somatic mutations uncovered by WES-tumor (Figure 2A). WES-cfDNA identified a variety of mutation types: frameshift (including insertions and deletions), missense, noncoding (splicing), and nonsense mutations. Additionally, WES-cfDNA discovered 14 high-confidence somatic mutations not identified by WES-tumor. Importantly, in the absence of the tumor specimen, WES-cfDNA could identify 68 of the 88 (77.3%) total mutations that could be identified by both techniques. The complete list of somatic mutations and all the sequencing parameters and genomic annotation are depicted in Table S1. Our results reveal the high capacity of tumor-free WES-cfDNA for global detection of somatic mutations, which was similar to that of tumor DNA sequencing. There was also high concordance in the identification of copy number variation (CNV) between WES-cfDNA and WES-tumor (Figure 2B).

#### *KDR* somatic mutations are recurrent in cancer

After the exclusion of the variants present also in the general noncancerous population, *KDR bona-fide* cancer mutations were found in 691 (1.6%) of the 33,320 cancer samples in the Catalogue Of Somatic Mutations In Cancer (COSMIC, *v80*) database. Furthermore, somatic mutations in *KDR* occurred in 481 (2.6%) of the total samples and in 56 (2.7%) of the 2,102 CRC samples from almost 19,000 cancer samples included in the AACR Project Genomics Evidence Neoplasia Information Exchange (GENIE) project (Figure 3A). We also analyzed *KDR* mutations from a third non-overlapping cancer genomic project, The Pancancer Analysis of Whole Genomes (PCAWG) study, which includes whole-genome sequencing data from 2,800 samples of 37 distinct types of cancer. From the 35 CRC samples included in the PCAWGS project, 4 (11.5%) harbored *KDR* mutations considered pathogenic. Although smaller than the other datasets, the PCAWGS database presents the advantage that whole-genome sequencing yields a more homogeneous coverage than whole-exome sequencing, enabling greatly more accurate copy number analysis results. Interestingly, 15 of the 35 (43%) CRC samples in PCAWGS had an amplification of the *KDR* gene. Moreover, *KDR* mutations were identified in 2.5% of the recently published Memorial Sloan Kettering Cancer Center (MSK-IMPACT) Clinical Sequencing cohort, including almost 11,000 samples from several cancer types^13^ (Figure 3A). The *KDR* somatic mutations found in the cancer genomics databases are shown in Table S2.

**Figure 3:**
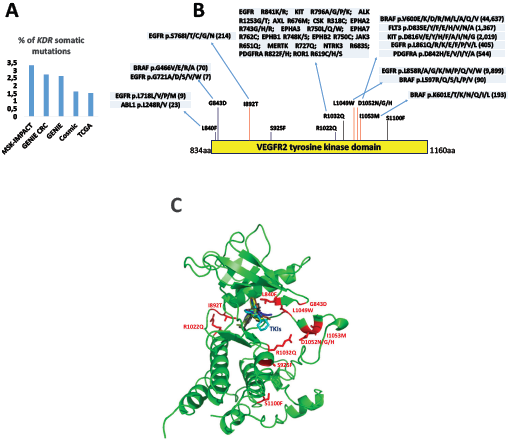
Recurrence of *KDR* /VEGFR2 oncogenic mutations in human cancers. A) Frequency of *KDR* somatic mutations in large cancer genomic sequencing projects including more than 70,000 cancer samples. Common germline polymorphisms (figure S26) were excluded from these analyses and only cancer-exclusive mutations were considered. B) Representative results from the search of genomics and protein databases, showing several mutations, structurally analogous to those of *KDR*, identified in other cancer-relevant kinases. C) 3D structural distribution of the VEGFR2 cancer mutants (red), and localization of TKI binding sites (blue).

Importantly, *KDR* was statistically classified as a *driver* of colorectal cancers (*p* = 2.0 × 10^-7^) and melanomas (*p* = 4.3 × 10^-5^) in the PCAWGS database, and a *driver* of lung adenocarcinoma and glioblastoma multiforme tumors by the Integrative Onco Genomics (IntOGene) platform^14^, which applies OncodriveFM, OncodriveCLUST, MutSigCV, and OncodriveROLE cancer driver detection methods.

#### Mutations structurally analogous to the recurrent *KDR* mutations occur in other kinases in cancer

In addition to the previously unrecognized occurrence of *KDR* mutations in cancer, we also found that structurally similar mutations occur in hot-spot residues of other kinases, clinically relevant for human cancers. For example, the analogous mutations of VEGFR2 L1049W (EGFR L858) and of VEGFR2 D1052N/G/H (FLT3 D835, KIT D816, EGFR L861, and PDGFRA D842) can be consider cancer hot-spots, as they are mutated in 58,871 cancer samples from the COSMIC database alone (Figures 3B-C).

Importantly, we found that R1032Q (COSM192176) was the most frequent mutation in three large non-overlapping cancer databases (TCGA, GENIE, PCAWGS), suggesting that R1032 is a possible mutational hot-spot in VEGFR2. Indeed, previous studies have shown that R is the most mutated residue in many kinases in different cancer types owing to the predominant C>T (G>A) transition mutability in CpG sites (four of the six codons for R have a CpG dinucleotide in the first and second positions)^15^. Moreover, our comparative structural analysis confirmed the mutation predisposition of this specific conserved R residue among several oncogenic kinases: VEGFR2 **R**1032Q; EGFR **R**841K/R; KIT **R**796A/G/P/K; ALK **R**1253G/T; AXL **R**676M; CSK **R**318C; EPHA2 **R**743G/H/R; EPHA3 **R**750L/Q/W; EPHA7 **R**762C; EPHB1 **R**748K/S; EPHB2 **R**750C; JAK3 **R**651Q; MERTK **R**727Q; NTRK3 **R**683S; PDGFRA **R**822F/H; ROR1 **R**619C/H/S (COSMIC and TCGA databases; Figures 3B and S9). In addition to R1032Q, identified in 20 cancer samples from the cancer sequencing projects analyzed, the R1032X nonsense mutation was detected in seven samples from distinct types of cancers, such as colorectal, lung, prostate, and haematopoietic cancers, as well as in melanomas and glioblastomas.

### FUNCTIONAL STUDIES OF THE VEGFR2 CANCER MUTANTS

#### L840F causes therapy resistance

In agreement with the phenotype we observed in the patient, the patient-derived Avatar model (Avatar^VEGFR2:L840F^) did not respond to multiple anti-VEGF and VEGFR2 inhibitors, whereas the Avatar^VEGFR2:WT^ model, used as a control, was sensitive to all the treatments (Figure 4A). In order to understand better how VEGFR2 L840F caused such broad *in vivo* resistance to anti-angiogenic drugs we conducted both *in silico* and biochemical analyses.

**Figure 4:**
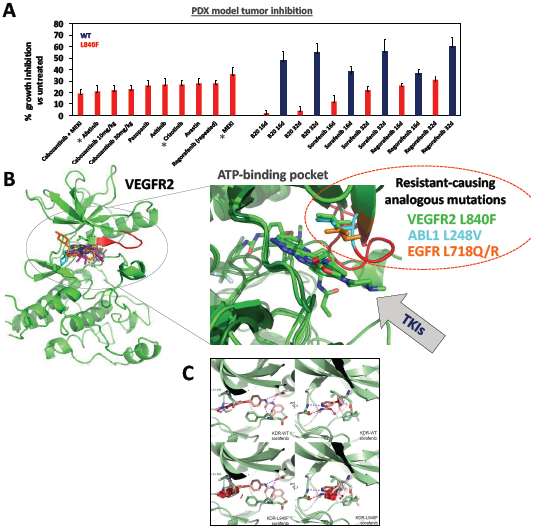
The VEGFR2 L840F mutant leads to broad and strong cancer therapy resistance. A) Growth inhibition of the patient-derived xenograft (PDX) Avatar model carrying the *KDR*/VEGFR2 L840F (red) after three weeks of treatment with anti-VEGF drugs (B20/murine and bevacizumab/human), VEGFR2 kinase inhibitors (axitinib, cabozantinib, cabozantinib:MEK inhibitor combo, pazopanib, regorafenib, sorafenib), or inhibitors of other kinases, such as afatinib (EGFR), crizotinib (MET), and MEK inhibitor (MEKi). A second CRC PDX model, carrying the *KDR*/VEGFR2 WT (blue), was used as a positive control, and was treated with some of the above drugs. To analyze the inhibition of tumor growth promoted by each tested drug, tumor volumes of the untreated mice were set as 100% growth and used as reference for the measurement of the treated animals (tumor volume of treated mice divided by tumor volume of untreated mice). B) Localization of structurally analogous L residue mutations in VEGFR2 (L840), EGFR (L718), and ABL1 (L248). A close-up view of the entrance of the ATP-binding pocket domain is shown in green for VEGFR2 L840F, in light blue for ABL1 L248V, and in orange for EGFR L718Q/R. Patients with these mutations are all refractory to treatment with tyrosine kinase inhibitors, which directly bind to these L residues (see also Figures S10-S20 and supplementary discussion for detailed information). C) Sorafenib binding to WT (upper panels) and L840F (lower panels) VEGFR2. Upper: Crystal structure of the WT VEGFR2 kinase domain bound to sorafenib (pdb accession code: 3WZE). VEGFR2 is shown in green and sorafenib in pink, with hydrogen bond interactions shown in magenta. Two views are shown, rotated 90° relative to each other. Lower: The L840F mutation is modelled in grey. Resulting clashes between F840 and sorafenib are shown as red discs.

Our 3D structural analysis revealed that L840 is located exactly at the entrance of the ATP-binding pocket of the tyrosine kinase domain of VEGFR2 (aa840-***<u>L</u>***GXGXXG-846aa), and it forms hydrophobic interactions with many FDA-approved small-molecule kinase inhibitors. Figures 4B and S10-S14 illustrate the direct contacts of VEGFR2 L840 with lenvatinib, sorafenib, axitinib and tivozanib, while the bindings of the VEGFR2 L840 analogous mutations ABL1 L248 with bosutinib, nilotinib, dasatinib, imatinib and axitinib, and EGFR L718 with WZ4002 are shown in Figures S15-S20.

We then modeled the L840F mutation *in silico* and observed that a clash occurs between the tyrosine kinase inhibitors (TKIs) and the F840 residue (Figure 4C), which would prevent the original mode of TKI binding to the receptor. Molecular dynamics simulations of the L840F mutant VEGFR2 also show that most of the conformations of F840 observed in the simulations are not compatible with inhibitor binding (Figures 4C, S21). Consistent with the computational model, *in vitro* kinase assays with recombinant VEGFR2 kinase domains showed that WT VEGFR2 has high kinase activity, whereas L840F VEGFR2 has impaired kinase activity, suggesting possible loss of ATP binding (Figure 5A). Consistent with these results, Y1175 phosphorylation of L840F VEGFR2 was significantly reduced compared to WT VEGFR2, both in human embryonic kidney (HEK293) cells transiently transfected with WT or L840F VEGFR2 and in porcine endothelial (PAE) cells stably expressing WT or L840F VEGFR2 (Figure S22). We also explored whether the molecular clash caused by the L840F mutation would prevent TKI inhibition *in vitro*. For this, we rescued L840F kinase activity by increasing the enzyme concentration a thousand times and evaluated its inhibition by TKIs. Indeed, we found that whereas WT VEGFR2 was sensitive to axitinib, cabozantinib, dovitinib, and levantinib, L840F VEGFR2 was resistant to all these drugs, albeit at distinct levels (Figure 5B).

**Figure 5:**
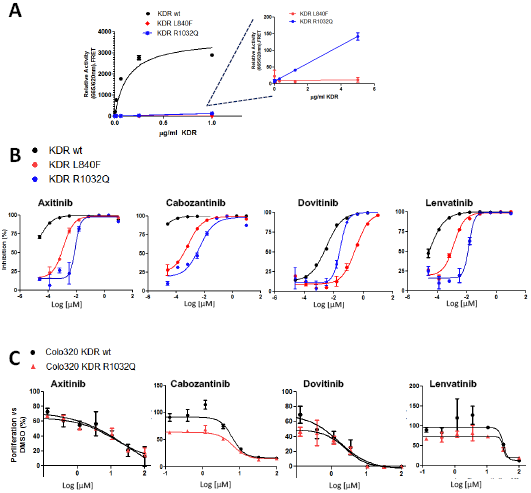
Biochemical and *in vivo* characterization of VEGFR2 L840F and R1032Q cancer mutants. A) Kinase activity assays of WT (black), L840F (red), and R1032Q (blue) VEGFR2 kinase domains. As expected, kinase activity was observed in the WT protein, whereas both VEGFR2 mutants were incapable of substrate phosphorylation, indicating that the mutations are kinase-inactivating. B) Inhibition of kinase activity of WT (black), L840F (red), and R1032Q (blue) VEGFR2 kinase domains by VEGFR2 inhibitors axitinib, cabozantinib, dovitinib, and lenvatinib. WT VEGFR2 kinase was sensitive to the four inhibitors, especially to axitinib and cabozantinib. The concentration of the mutants was increased 1,000 times to achieve a measurable kinase activity, which was consequently measured in the presence of TKIs. C) Proliferation assays of Colo320 cell lines, stably expressing WT and R1032Q VEGFR2, upon treatment with TKIs. The expression of the VEGFR2 R1032Q hot-spot mutant in Colo320 cell lines (WT to *KRAS*/*NRAS*/*BRAF*/*PIK3CA* and mutated to *TP53* and *APC*) increased sensitivity to cabozantinib. Consistent with these results, the MDST8 CRC cell line, constitutively expressing R1032Q VEGFR2, was also more sensitive to cabozantinib (Figure S23).

#### R1032Q confers sensitivity to strong VEGFR2 inhibitors

In addition to L840F VEGFR2, we characterized the kinase activity of R1032Q VEGFR2 with our *in vitro* kinase assays, and we found that similar to the L840F, the R1032Q mutation greatly reduced VEGFR2 kinase activity (Figure 5A), although the structural features underlying this phenotype would be distinct. Whereas L840F most likely blocks ATP and TKI entrance to the ATP-binding site leading to kinase inactivation and TKI resistance, R1032Q does not directly interfere with the ATP-binding pocket. Instead, R1032Q affects the kinase catalytic motif DxxxxN (aa1028-Dxxx**<u>R</u>**N-1033aa), which is present in all known protein kinases and is directly involved in the catalytic mechanism of the enzyme^16^. Notably, *in vitro* analyses by different research groups have identified the following mutations in R1032-analogous residues of other kinases that greatly impair kinase activity: EGFR R841^17-19^, c-KIT R796^20^, ALK R1253^21^, CSK R318^22^.

Since R1032, contrary to L840, does not directly participate in receptor:TKI binding (Figures S10-20), we asked whether R1032Q VEGFR2 would be inhibited by TKIs. *In vitro* kinase assays showed increased sensitivity of R1032Q VEGFR2 to TKIs (Figure 5B). Furthermore, proliferation studies with the Colo-320 colorectal cell line, which has a similar mutation profile as that of the patient’s tumor (WT *KRAS*/*NRAS*/*BRAF*/*PIK3CA* status, and *TP53* and *APC* mutations), showed that stable expression of R1032Q VEGFR2 conferred sensitivity to lenvatinib (growth inhibition (GI_50_) = 20.8 for the R1032Q compared to 36.4 for WT VEGFR2) and to cabozantinib (GI_50_ = 2.5 for the R1032Q compared to 7.9 for WT VEGFR2) (Figure 5C). Moreover, cabozantinib treatment of the MDST8 CRC cell line, naturally harboring the *KDR* R1032Q mutation, led to a prominent decrease in cell growth rate *in vitro* and diminished the high constitutive ERK phosphorylation levels (Figure S23). Importantly, we found that such downstream inhibition was specific to cabozantinib, a very strong VEGFR2 (0.035 nM) and c-MET (1.3 nM) inhibitor, and occurred in cells treated in the absence or presence of VEGF (Figure S23).

#### Oncogenic effects of L840F, R1032Q, and other VEGFR2 cancer-related mutants

Based on recent studies that comprehensively showed that Protein Kinase C β (PKCβ) and Mixed-Lineage Kinase 4 (MLK4) loss-of-function mutations play an oncogenic role in CRC^23,24^, we asked if in addition to modulating the response to TKIs, VEGFR2 cancer mutants could promote tumor growth. Indeed, we showed that Colo-320 cells stably expressing L840F VEGFR2, even when injected in small numbers and without matrigel, could generate tumors that reached the established humane endpoint (Figure 6). In addition, the R1032Q hot-spot mutant, as well as other cancer-related VEGFR2 mutants (D717V, G800D, G800R, G843D, S925F, R1022Q, R1032Q, and S1100F), promoted tumor growth *in vivo*. Importantly, Colo-320 expressing empty vector (EV) or the kinase inactive dominant-negative K868M VEGFR2 did not generate tumors within 120 days after cell injections. Our data suggest that cancer-associated VEGFR2 mutants might have oncogenic potential (Figure 6).

**Figure 6:**
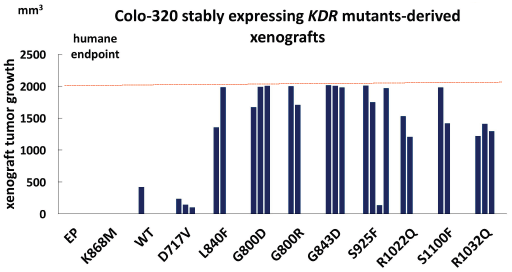
Oncogenic potential of VEGFR2 cancer mutants in xenograft assays. Colo320 CRC cell lines were used for xenograft studies, as they resemble the mutation profiling of the studied patient’s tumor, with wild-type *KRAS*/*NRAS*/*BRAF*/*PIK3CA* status, and *TP53* and *APC* mutations. Colo320 cells stably expressing different VEGFR2 mutants were injected subcutaneously in four immune deficient mice (each blue bar represents one mouse). Results of xenograft growth after two months following injections or when tumors reached the established humane endpoint are shown.

## DISCUSSION

Our study brings three main novelties to the field of cancer translational research. First, we established the potential of WES-cfDNA as a global tumor-free genomic platform to explore genetic causes of primary resistance to cancer therapies. Second, following the identification of the L840F VEGFR2 clonal, somatic mutation in a highly refractory mCRC patient by WES-cfDNA, we performed functional analyses that provide a mechanistic understanding of *KDR*/VEGFR2 somatic mutations as genetic modulators of the response to anti-angiogenic drugs. Lastly, we explored genomic sequencing databases and showed that recurrent *KDR*/VEGFR2 somatic mutations, which are structurally analogous to cancer host-spot mutations in other kinases, occur in 1-3% of human cancers.

The high capacity of WES-cfDNA for portraying the somatic mutation and copy number variation landscapes of tumors has immense potential for research in translational oncology^12,25,26^. During the last decade, cancer genomics projects proved whole-exome sequencing of tumor samples, WES-tumor, to be a powerful and affordable technique that is still considered the gold-standard platform for the unbiased discovery of somatic mutations and the characterization of cancer genome landscapes. A key limitation of WES-tumor is the use of tumor fragment(s) acquired during surgery or biopsy as the source of cancer DNA. The future implementation of WES-cfDNA analysis into genomic projects will enable the investigation of a much larger number of cancer patients, including those for whom only blood/plasma samples are available because of an inoperable tumor or because the tumor localization renders biopsy unsafe. Additionally, WES-cfDNA can uncover somatic mutations not represented by the WES-tumor, therefore complementing the knowledge about the patients’ cancer and allowing the identification of more drug targets and therapeutic options. Following these promising results, we have implemented the WES-cfDNA prospective gene discovery and clonal evolution platform into the NCT02795650 ongoing clinical trial and have identified potentially therapeutic relevant mutations in *BRCA2* and *ROS1* in advanced pancreatic cancer patients (Toledo RA *et al*, manuscript in preparation). VEGFR2, EGFR, ABL1, and PDGFRA are major cancer therapeutic targets with several TKIs already approved in the clinic. However, whereas TKI sensitizing and resistant mutations in EGFR, ABL1, and PDGFRA have already been well characterized, much less is known about mutations in VEGFR2 and their clinical implications (supplementary discussion, points I and II). The main reasons for this surprisingly limited amount of information on *KDR*/VEGFR2 mutations and their pharmacological impact are: a) the concept that VEGFR2 is mostly expressed and playing a role on endothelial but not on tumor cells, and b) the previous notion of paucity or even absence of *KDR* mutations in cancers, as suggested by small genetic screening studies^27^. Thus, after the identification and confirmation of the L840F clonal, somatic mutation in a refractory mCRC cancer (supplementary discussion, point III), we aimed to further characterize the frequency and the role of *KDR*/VEGFR2 mutations across human cancer. To this end, we performed an expanded screening of cancer genome sequencing projects and carried out a series of comprehensive *in vitro* and *in vivo* functional experiments.

Our experiments with cell lines, animal models, and biochemical assays showed that the L840F ATP-binding pocket domain mutation causes very strong and broad resistance to anti-VEGF and VEGFR2 inhibitors (supplementary discussion, point V). On the contrary, we showed that the R1032Q kinase hot-spot mutation is sensitive to the strong VEGFR2 inhibitor cabozantinib. These new findings demonstrate that, as occurring in EGFR, ABL1, and PDGFRA, mutations in VEGFR2 can have functional consequences and influence the efficiency of cancer targeted therapies. In addition, taking advantage of publicly available sequencing data from the TCGA, GENIE, PCAWGS, and MSK-IMPACT cancer genomic databases, we found that 1-3% of the 73,389 cancer samples analyzed harbored a potentially pathogenic somatic mutation in *KDR*/VEGFR2.

In agreement with our findings of a consistent occurrence of *KDR* mutations in cancers, three brief case reports describing patients with a *KDR*-mutated tumor became very recently available. Although no accompanying experimental data were provided, the patients were treated with anti-angiogenic drugs based on the expectation that the VEGFR2 mutant could be a potentially sensitive target, as follows. Knepper *et al.* reported a prolonged complete response to pazopanib in a metastatic basal cellular carcinoma patient carrying the somatic *KDR*/VEGFR2 R1032Q^28^. This case represents the clinical confirmation of our preclinical experimental results, showing that R1032Q is a sensitizing mutation for at least some strong VEGFR2 inhibitors. The second case was a mCRC patient treated with 5-fluorouracil-bevacizumab for six cycles with progression of the disease. A panel of 47 genes was analyzed by NGS and in addition to common mutations in *APC* and *KRAS*, the *KDR* R961W mutation was observed in nearly 30% of the reads^29^. The patient was offered low dose regorafenib (80 mg/day) that promptly needed to be reduced to 40 mg/day owing to secondary effects. After 3 months of treatment, imaging scans reveled remarkable improvement in the hepatic metastases, abdominal and retroperitoneal lymph nodes, and rectosigmoid colon hypermetabolic lesions. The third case was a mCRC patient who progressed to FOLFOX-cetuximab, folinic acid/fluorouracil-cetuximab, and was then treated with successive irinotecan-cetuximab plus ramucirumab^30^. After two cycles, the patient developed a sporadic expanding angioma that was shown by WES to be clonal and carry the *KDR* T771R as the only somatic mutation. The authors suggest that *KDR* T771R may offer a proliferative advantage in the setting of ramucirumab treatment. A fourth *KDR*-mutated case was recently identified during the generation of tumor organoids in a series of twenty consecutive CRC patients, however no information on therapeutics was provided (supplementary discussion, point VI),

Collectively, the data from these three cases show no evidence of benefit by indirect or direct blocking of the extracellular domain of VEGFR2 mutants (i.e. with bevacizumab and ramucirumab). Interestingly, the two patients with tumors carrying mutations located in the kinase domain (R1032Q and R961W) responded well to VEGFR2 inhibitors (pazopanib and regorafenib), whereas the other two patients with tumors carrying mutations outside of the kinase domain (T771R and L840F) did not respond to TKIs. These latter patients not only did not respond, but it seems that blocking of VEGFR2 had adverse effects. For example, our mCRC patient carrying the L840F ATP-binding domain mutation progressed very fast to regorafenib (Figure 1A) and the mCRC *KDR* T771R-mutated patient developed an expanding angioma after ramucirumab^30^.

The classical understanding of kinases as straightforward “activators/phosphorylators” of downstream pathways has been greatly expanded and at least six types of cancer kinase mutations have been discovered and characterized, including genesis/extinction of phosphorylation sites, node activation/inactivation, and downstream/upstream rewiring^31^. The L840F and R1032Q VEGFR2 mutations impairing kinase activity would best fit the upstream rewiring mutant kinase oncogenic model, whereby the mutant receptor recruits unusual partner(s), which are responsible for MAPK pathway activation (Figure S24). This has been demonstrated for BRAF impaired mutants that recruit CRAF, promoting alternative MAPK pathway activation and leading to BRAF inhibitor therapy resistance. Interestingly, this BRAF:CRAF pathway rewiring confers *de novo* clinical sensitivity to dasatinib (ABL, SRC, and c-Kit inhibitor)^32^. In our study, tumor growth of the Avatar^VEGFR2:L840F^ model was not significantly reduced upon treatment with inhibitors of EGFR (afatinib), MET (crizotinib), and MAPK (MEKi) (Figure 4A), and the possible rewiring partner of L840F VEGFR remains unknown (supplementary discussion, point VI). Our findings resonate the difficulties of therapeutically targeting kinases with mutations in the L residue located at the entrance of ATP/TKI binding.

The current study is the first to implicate in a robust manner VEGFR2 as a direct regulator of the efficacy of anti-angiogenic therapies in common cancers and can facilitate answering a long-standing unmet medical question regarding biomarkers of response to anti-angiogenic drugs. A complete functional and clinical characterization of *KDR* cancer mutations will be necessary to classify phenotypes of cancers carrying VEGFR2 mutants as neutral, sensitizing, or resistant to anti-VEGF and VEGFR2 inhibitor treatments. These data should be taken into consideration for the future design of small population *basket* clinical trials based on *KDR* genotypes. Moreover, a retrospective or prospective investigation of *KDR* gene expression within CRC cohorts treated with anti-angiogenic drugs could also be of interest, since we showed that among more than 1,000 patients analyzed, those with mCRC with high tumor *KDR* gene-expression did significantly worse (a finding confirmed in four independent CRC cohorts, Figure S25).

## Author contributions

Conceived and oversaw the project: RAT, JLMT, MH; conceived the plasma cfDNA whole-exome sequencing gene discovery strategy: RAT; diagnosed and treated the patient: EG, EV, RA, AC, MH; provided clinical and sample management: FS, SP; designed, performed, interpreted experiments: RAT, MM, JM, MC, TSP, MIA, AO, ADM, DL, CBA, OD, JLMT; performed in vivo studies: NB, YD, VB; performed protein structural analyses: TP, MC, DL; performed genomic and protein database analyses: TP, RAT; wrote the paper: RAT with inputs from JLMT, EG, MM, MH and all authors.

## Acknowledgments

We are grateful to Dr. Pedro P. López-Casas and Manuel Muñoz (CNIO Gastrointestinal Cancer Unit) for their valuable technical and administrative assistance; to Dr. Javier Muñoz and Eduardo Zarzuela (CNIO Proteomics Core Unit) for assistance with protein mutation analysis; to Dr. Diego Megías and Manuel Pérez (CNIO Confocal Microscopy Core Unit) for their assistance with the tissue immunofluorescence preparation; to Prof. Patricia L. Dahia (Department of Medicine of the University of Texas Health Science Center at San Antonio) and for Dr. Rodrigo Dienstmann (Vall d’Hebron Institute of Oncology, Barcelona, Spain) for her critical reading and suggestions on the manuscript; to Kurt Ballmer-Hofer (Paul Scherrer Institute, Switzerland) for his involvement at the beginning of the functional *in vitro* experiments. RA Toledo was a recipient of a research fellowship from the National Council for Scientific and Technological Development (CNPq). The authors would like to thank especially the patient and his family for their participation in the study.

## Supplementary discussion, #1-6

### #1) The timing of occurrence of the KDR/VEGFR2 L840F mutation

The time point when kinase mutations occur in cancer is still under debate, with different studies showing different results. Independent N-ethyl-N-nitrosourea (ENU)-induced cell-based mutagenesis studies identified the ABL1-L248 and EGFR-L718 mutations (analogous to VEGFR2-L840, which we found in the highly refractory mCRC patient) as a recurrent mechanism of acquired resistance to ABL112 and EGFR13 inhibitors, suggesting that these mutations are secondary events emerging after therapies with kinase inhibitors. Accordingly, the same leucine mutations were observed in patients that acquired clinical resistance to the same ABL1 and EGFR kinase inhibitors tested *in vitro*^1,2^. However, other studies have shown the existence of resistance-associated kinase mutations before treatment^3,4^.

Our finding of the *KDR*/VEGFR2 L840F mutation in the plasma cfDNA collected before frontline treatment with FOLFIRI-cetuximab shows that KDR mutations can occur independently of pressure from therapies inhibiting the VEGF/VEGFR pathway (Figure 1, S1). Importantly, the mutated allele frequencies (MAFs) of *KDR*/VEGFR2 L840F were similar to the MAFs of trunk CRC mutations, such as those of APC and TP53. The MAF of *KDR*/VEGFR2 L840F in the tumor sample was approximately 30%, while the MAFs for both the APC p.E1322EX and TP53 p.Y220SY mutations were approximately 50%. Importantly, the MAF of *KDR*/VEGFR2 L840F in plasma (11%) was higher than that of APC (8%), whereas the MAF of TP53 was the highest (18%). These data support the notion that the *KDR*/VEGFR2 L840F mutation occurred independently of therapeutic pressure and provide strong evidence that it was a clonal event occurring concomitantly or somewhat later than the *APC* and *TP53* mutations.

### #2 *KDR* /VEGFR2 mutations in angiosarcomas

Previous gene screening studies investigated a limited number of cancer samples and failed to identify *KDR* mutations. The only tumors, where *KDR* somatic mutations have been better characterized, were angiosarcomas^5^. Few *KDR* mutations spanning the extracellular (T717V), transmembrane (T771R), and kinase (A1065T) domains of VEGFR2 were initially identified in a specific subset of patients (cutaneous angiosarcomas of the breast)^5^. Importantly, the authors conducted *in vitro* experiments that showed that the T717V and A1065T VEGFR2 mutants were sensitive to sorafenib and sunitinib, whereas no data were shown for the T771R variant. The same year that these findings were published, the efficacy of sorafenib for the treatment of angiosarcoma patients was evaluated in the context of two phase-II trials. These clinical studies showed limited^(6)^ or no clinical response^7^. Importantly, the former trial screened the tumors of all referred patients for *KDR* mutations, but none was identified, and the study failed to show a benefit of sorafenib in the treatment of the enrolled *KDR*-WT angiosarcoma patients^7^. Although the scarcity of *KDR* mutations in angiosarcoma patients has been confirmed in other cohorts^8^, a recent study, focusing exclusively on angiosarcomas of the breast, reported up to 26% of KDR mutations in the T771 residue (mainly T771R, but also T771K) located at the transmembrane domain of VEGFR2^9^.

The finding of VEGFR T771 variants in angiosarcomas of the breast by two groups^5,9^ could suggest implication of these variants in the disease. However, so far there are no experimental data indicating a potential functional role of T771R and T771K. Importantly, the evolutionary non-conservation of the T771 residue is a strong argument against the pathogenicity caused by T771 variants. For example, the missense variant T771M is a known benign germline polymorphism segregating in Latino, European, and mainly in African populations (rs149745504, allele frequency^10^: afr: 0.42%, eur: < 0.1%).

### #3 VEGFR L840F-analogous mutants occurring in the L residue at the entrance of the ATP-binding pocket in other kinases lead to broad and strong resistance to kinase inhibitors

EGFR L718 and ABL1 L248 are structurally analogous to VEGFR2 L840. Mutations in EGFR L718 (L718M/P/V) have been previously identified in lung cancers^2^, and very recently the EGFR L718Q mutation was identified as the most frequent cause of resistance to the EGFR inhibitor WZ4002^(11)^. EGFR L718Q also causes resistance to CO-1686 and AZD9291^11,12^, clearly pointing L718 as a key residue, especially implicated in TKI treatment efficacy. Importantly, L718 is also indirectly involved in the broad resistance to most clinically available EGFR inhibitors caused by the EGFR T790M hot-spot mutation. EGFR T790M causes structural changes that narrow the hydrophobic slot formed between L718 and G796^13,14^.

Consistent with the hypothesis that a highly resistant phenotype correlates with mutations in the entrance of ATP-binding pocket, the ABL1 L248 residue has also been implicated in TKI resistance. ABL1 L248 mutations were found in imatinib-resistant CML patients^15^, and *in vitro* studies revealed a highly resistant phenotype of the ABL1 L248V mutant to imatinib (IC_50_: 2100 nM vs 260 nM for the WT, and nilotinib (IC_50_: 100 nM vs 13 nM for the WT)^16^.

Importantly, mutations in these L residues also seem to negatively interfere with important opportunities for clinical drug-repositioning. It was recently shown that the ABL1 T315I hot-spot gatekeeper mutant is effectively inhibited by axitinib (a VEGFR2 inhibitor); however, the three ABL1 mutations involving the L248 residue (L248R+F359I, L248V, L248R) promoted the highest levels of *in vitro* refractoriness among the 28 different ABL1 mutations investigated^17^.

These data are consistent with our finding of the VEGFR2 L840F mutation in a highly refractory mCRC patient, and reinforce the concept that somatic kinase mutations involving the L residue at the entrance of the kinase ATP-binding pocket (LGXGXXG) cause broad and strong TKI resistance, and are very difficult to target therapeutically. To our knowledge, the possible existence of escape molecular mechanisms of ABL1 L248 and EGFR L718 mutants that could be therapeutically targeted remains elusive and no effective treatment has been discovered for patients with tumors carrying these mutations. We found similar results with the VEGFR L840F, as described below.

### #4 Genomic findings of new KDR somatic/tumor-exclusive/non-germline-polymorphisms in cancer samples

It is important that the new genomic findings of the current study, demonstrating the occurrence of rare but recurrent somatic *KDR*/VEGFR2 mutations across common cancers, should be considered separately from studies correlating angiogenic responses with VEGF/VEGFR germline polymorphisms^(18,19,20)^, as well as from the published genetic screening studies that inaccurately reported the two VEGFR2 variants V297I and Q472H as likely pathogenic/cancer-driven *KDR* mutations^(21,22)^. V297I (rs2305948) and Q472H (rs1870377) are not cancer-exclusive mutations but non-pathogenic germline polymorphisms with allele frequencies between 10% and 20% across non-cancerous-enriched populations from the 10,000 genomes^(10)^ and EXAC^(23)^ databases (Figure S26). In addition to somatic mutations, we found that *KDR* copy number gain occurs in more than 40% of the CRC samples included in the PCAWGS database (analyzed by whole-exome sequencing). Together with previous studies that have shown frequent gain of the *KDR* locus in non-small-cell carcinomas (approximately 30%)^(24)^, these findings indicate that somatic *KDR* gene gain is a frequent somatic event in common cancers.

### #5 The *KDR* somatic mutation G843D, analogous to the oncogenic hot-spot *KRAS* G12D, was identified in primary mCRC and the corresponding organoid model

Very recently, a library of tumor organoids derived from a series of twenty consecutive CRC patients was generated and shown to recapitulate the mutations and copy number variation profiles of the tumors^(25)^.

Interestingly, one of the patients (#P19) carried two *APC* WT primary tumors separated by > 10 cm that shared the same *TP53* and *BRAF* mutations but harbored different *KDR* mutations: the VEGFR2 E980Q and G843D mutants were identified in tumors P19a and P19b, respectively, as well as in their corresponding organoids. Interestingly, similar to the L840F mutation (***<u>L</u>***GXGXXG), the G843D mutation is located at the highly conserved ATP-binding pocket domain (LGX***<u>G</u>***XXG). Importantly, the VEGFR2 G843D mutant is analogous to the *KRAS* G12D mutant (COSM521), which is one of the most frequent hot-spot mutations in human cancers, and is highly oncogenic. KRAS G12D was found in all 162 cancer sequencing projects of *The Cancer Genome Atlas* (TCGA) project and in more than 14,000 cancer samples included in the COSMIC cancer database. Similarly, we have shown here that a colorectal cell line stably expressing the VEGFR2 G843D mutant promotes the growth of xenograft tumors (Figure 3D).

### #6 Exploring possible VEGFR2 L840F escape mechanism pathways

Decades of study of cancer kinases have taught us that mutations even in adjacent residues can have distinct effects on the kinase affinity for ATP binding and can lead to opposite phenotypes^16^. For example, as already mentioned, the L718R/V mutations in the ATP-binding domain of EGFR (LGXGXXG, structurally analogous to KDR/VEGFR2L840F) cause strong resistance to EGFR TKIs^(3)^, whereas adjacent G719S/A/C EGFR mutations (L***G***XGXXG) increase sensitivity to the same drugs^(11,26,27,28)^.

A similar case of mutation specificity is also found in the *BRAF* oncogene where the G466V/E (LGX***G***XXG) and G469A/V/E/R (LGXGXX***G***) ATP-binding pocket domain mutations cause opposite phenotypes: inactivating and activating, respectively. These findings have therapeutic implications because in contrast to the activating *BRAF* mutations, such as the classical *BRAF* V600E, the *BRAF* inactivating mutations confer *de novo* clinical sensitivity to dasatinib (ABL, SRC, and c-Kit inhibitor)^(29)^.

Based on these and other examples, we asked whether other molecular pathways could be indirectly implicated in the tumor growth and therapy resistance in the context of the VEGFR2 L840F mutation. For that, we treated the Avatar^VEGFR2:L840F^ model with EGFR (afatinib), MET (crizotinib), and MAPK (MEKi) inhibitors; however, none of these treatments reduced tumor growth significantly (Figure 4A), and the possible rewiring partner of the VEGFR L840F mutant that would explain the high levels of p-ERK and p-AKT observed in the Avatar^VEGFR2:L840F^ model remains unknown. Our findings resonate the difficulties of therapeutically targeting cancer mutations in the L residue located at the entrance of the ATP-binding pocket of the kinase domain.

## Supplementary Figures

**Figure S1:**
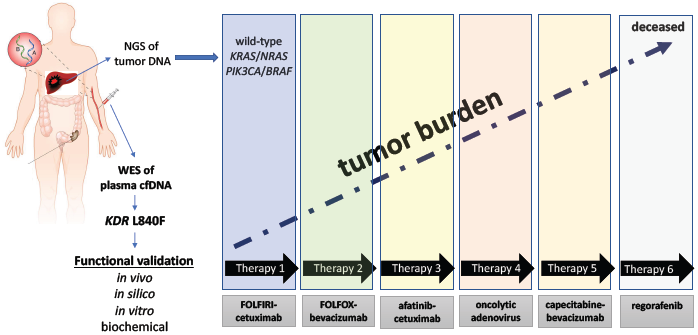
Outline of the genomic, tumor-free, resistance-associated gene discovery platform. Despite harboring no mutation in the known resistance genes (*KRAS, NRAS, BRAF, PIK3CA*) the metastatic colorectal cancer patient was refractory to six different standard and experimental therapies under development in the context of phase-I clinical trials. WES of plasma cfDNA identified the somatic L840F KDR/VEGFR, which was functionally validated *in vitro* and *in vivo*.

**Figures S2-S8:**
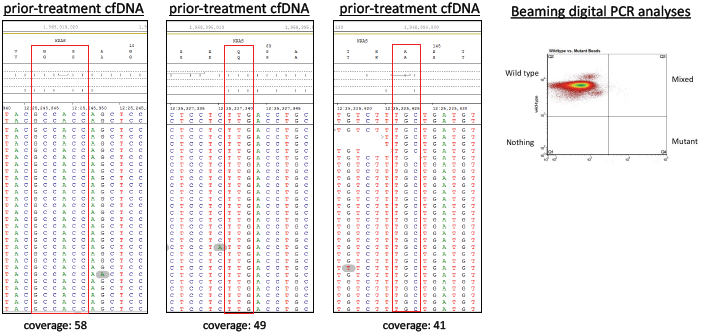
Read alignments from WES-cfDNA, showing no therapy resistance-associated hot-spot mutations in *KRAS* (Figure S2), *NRAS* (Figure S3), *BRAF* (Figure S3), and *PIK3CA* (Figure S4). Read alignments from WES-cfDNA and WES-gDNA (leukocytes), showing somatic mutations in *APC* (Figure S5), *TP53* (Figure S6), and *KDR* (S7). The *KDR*/VEGFR2 L840F mutation was confirmed to occur exclusively on the tumor cells by WES (Figure S7), Sanger sequencing (Figure S8), and TaqMan genotyping assay (Figure 1B). The right panel in Figure S2 shows the WT results by BEAMing (beads, emulsion, amplification, magnetics), an ultra-sensitive mutation detection technique that combines emulsion digital PCR and flow cytometry.

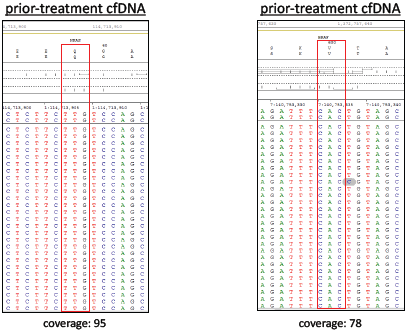

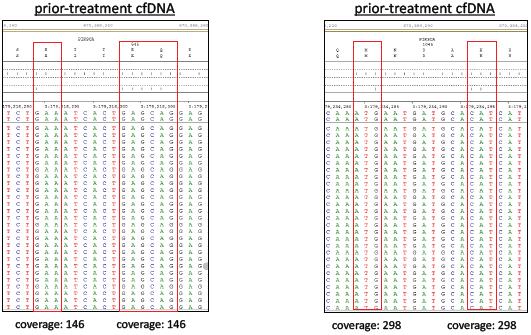

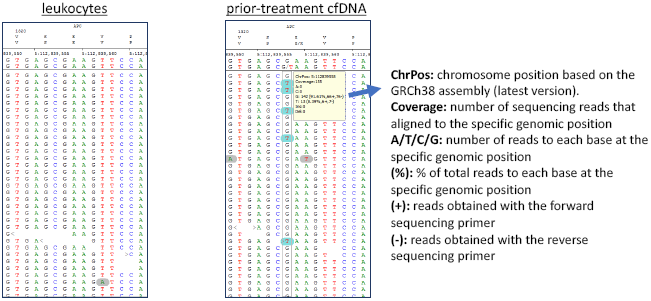

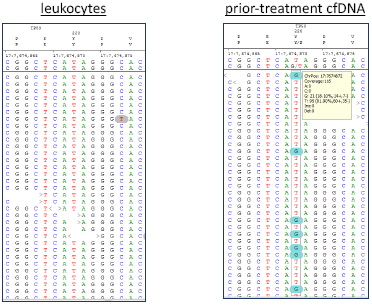

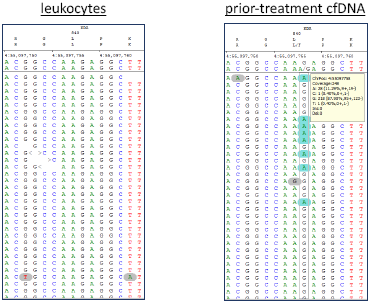

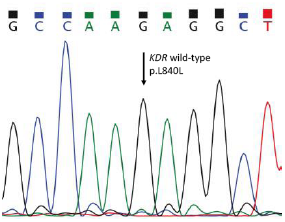

**Figure S9:**
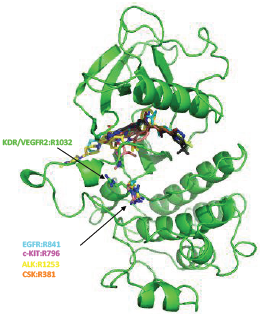
3D structure of the VEGFR2 receptor showing the localization of the R1032Q mutation, which is a VEGFR2 hot-spot mutation in human cancers, and R1032Q-analogous cancer mutations occurring in ALK, c-KIT, EGFR, and CSK. Several other R1032Q-analogous cancer mutations were retrieved from the COSMIC and TCGA databases, with a similar structural localization: AXL R676M; CSK R318C; EPHA2 R743G/H/R; EPHA3 R750L/Q/W; EPHA7 R762C; EPHB1 R748K/S; EPHB2 R750C; JAK3 R651Q; MERTK R727Q; NTRK3 R683S; PDGFRA R822F/H; ROR1 R619C/H/S.

**Figures S10-S20:**
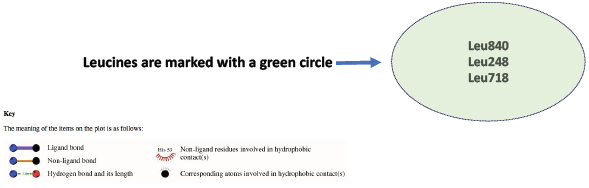
Analyses of interactions between tyrosine kinase receptors and kinase inhibitors, performed with LigProt. These analyses showed that VEGFR2 L840, ABL1 L248, and EGFR L718 directly bind to their corresponding kinase inhibitors (interaction marked with a green circle). On the contrary, the kinase inhibitors do not bind to VEGFR2 R1032Q or the analogous R residues in ABL1 and EGFR receptors.

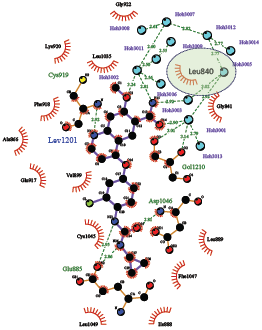

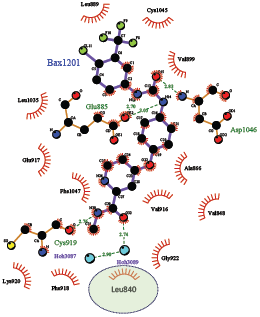

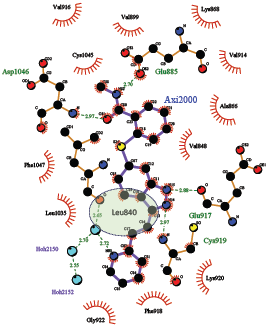

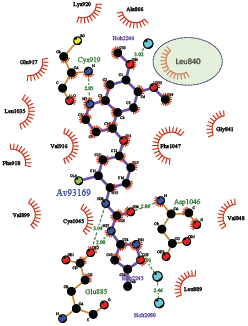

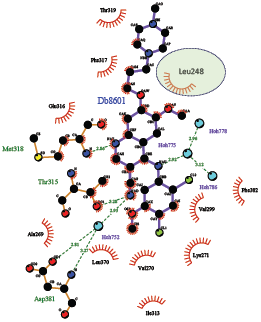

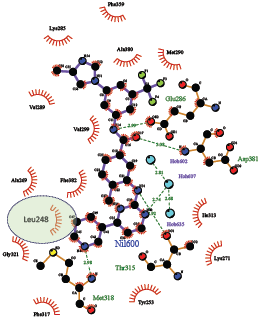

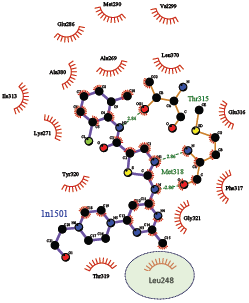

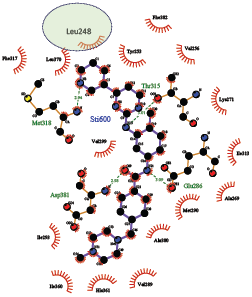

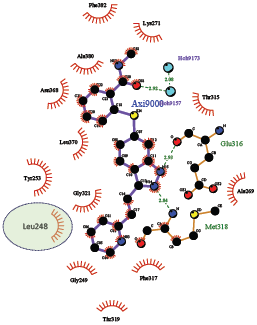

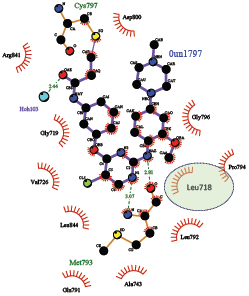

**Figure S21:**
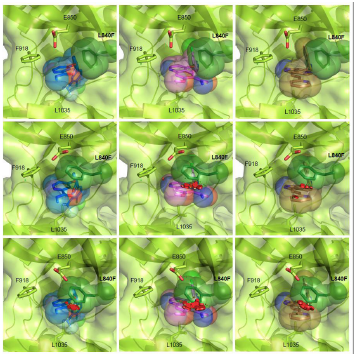
Analysis of TKI binding to alternative F840 conformations. Cluster analysis of molecular dynamics simulations with L840F VEGFR2 indicates that most of the adopted F840 conformations during the simulation are not compatible with inhibitor binding. Examples of conformations that can accommodate most TKIs (upper panels), some TKIs (middle panels) or no TKIs (lower panels) are shown. VEGFR2 is shown in green with the L840F mutation in darker green, sorafenib in blue, lenvatinib in magenta, and axitinib in brown. Clashes between VEGFR2 and the TKIs area are shown as red discs.

**Figure S22:**
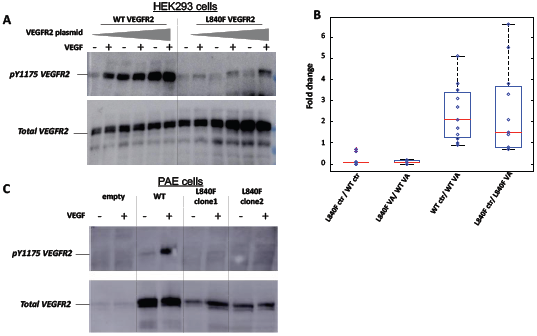
Effects of the L840F mutation on Y1175 VEGFR2 phosphorylation. (A) HEK293 cells were transiently transfected with increasing levels of a plasmid encoding for WT or L840F VEGFR2. Cells at 70% confluence were starved in 1% BSA/DMEM for 4 h, and then incubated in the absence or presence of 60 ng/ml VEGF_165_ for 10 min at 37 °C. Whole cell lysates were analyzed by western blotting, using antibodies against phosphoY1175 and total VEGFR2. Representative results are shown. (B) Quantification of 11 independent transient transfection experiments, as shown in (A). The intensity of the phosphoY1175 band was normalized against that of the total VEGFR2 band for all 4 samples, blotted in the same membrane: WT VEGFR2 in the absence (WT ctr) or presence (WT VA) of VEGF, and L840F VEGFR2 in the absence (L840F ctr) or presence (L840F VA) of VEGF_165_. Although the responsiveness to VEGF_165_ was similar for both WT and L840F VEGFR2 (WT VA / WT ctr and L840F VA / L840F ctr), the basal, as well as the VEGF-activated, levels of Y1175 phosphorylation were significantly decreased for L840F compared to WT VEGFR2 (L840F ctr / WT ctr and L840F VA / WT VA, respectively), suggesting that the L840F mutation impairs VEGFR2 kinase activity. (C) PAE cells stably expressing WT or L840F VEGFR2 were generated from a PAE cell line that does not normally express VEGFR2 (empty). Cells at 70% confluence were starved in 1% BSA/DMEM for 4h, and then incubated in the absence or presence of 60 ng/ml VEGF_165_ for 10 min at 37 °C. Whole cell lysates were then analyzed by western blotting, using antibodies against phosphoY1175 and total VEGFR2. Representative results are shown, highlighting the significant decrease in VEGF-induced Y1175 VEGFR2 phosphorylation in the presence of the L840F mutation.

**Figure S23:**
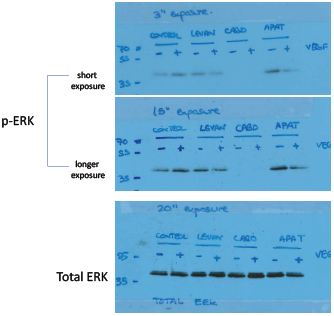
Levels of phosphorylated ERK following treatment of the MDST8 colorectal cell line, expressing R1032Q VEGFR2, with different kinase inhibitors in the presence or absence of VEGF. Cabozantinib specifically led to a strong and VEGF-independent inhibition of the MAPK pathway.

**Figure S24:**
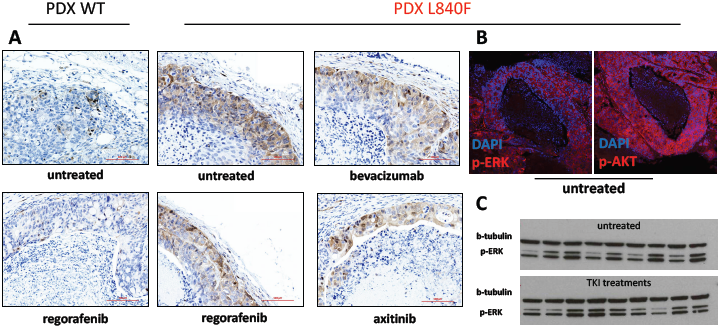
Assessment of the expression of phosphorylated ERK (p-ERK) in the WT and L840F VEGFR2 PDX Avatar models by immunohistochemistry (A), confocal microscopy (B), and immunoblotting (C). The MAPK pathway was activated in the L840F VEGFR2 mutant model and was not decreased following treatment with TKIs.

**Figure S25:**
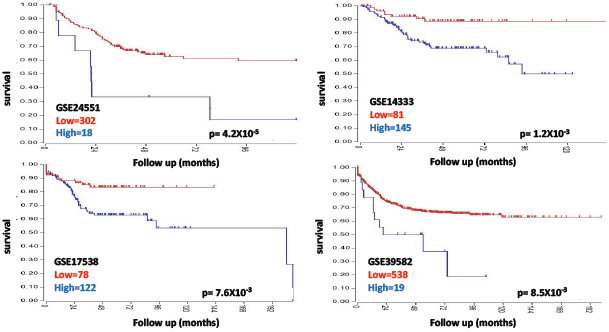
Overall survival according to *KDR* gene expression in four independent populations of CRC patients. Patients with low (red line) and high (blue line) *KDR* gene expression are shown. The number of patients in each group and the p value of the Kaplan-Meier estimate are shown.

**Figure S26:**
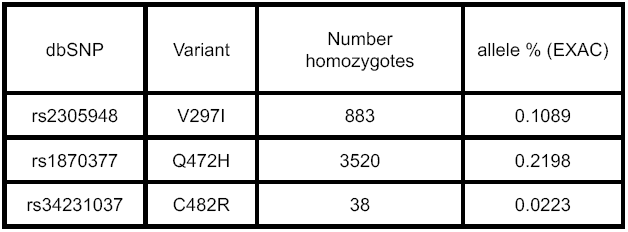
The three most common germline missense polymorphisms of *KDR* are shown. A large number of studies unequivocally describe these variants as pathogenic. We excluded these polymorphisms from all our analyses.

## ONLINE METHODS

### Study supervision

The study was approved by the institutional Review Boards of Hospital Universitario HM Sanchinarro and conducted in agreement with the Declaration of Helsinki and the International Conference on Harmonization of Good Clinical Practice guidelines. The patient gave written informed consent to participate in the study. Mice used in this research were treated humanely according to the regulations laid down by the Spanish National Cancer Research Centre (CNIO) Bioethics Committee.

### DNA extraction

DNA was extracted from leukocytes (gDNA), liver metastasis (tDNA), and basal and on-treatment plasma samples (cfDNA), using commercial kits according to the manufacturer’s instructions (Qiagen, Germany). The DNA amount was quantified with a Qubit™ Fluorometer (Thermofisher, USA) and reported in ng. cfDNA samples were also quantified using a modified version of human LINE-1-based quantitative real-time PCR and reported in genome equivalents (GE; GE being one haploid human genome weighing 3.3 pg). gDNA and tDNA was sheared to 300-bp fragments on a Covaris instrument (Covaris, Woburn, MA) according to standard procedures. The 2100 Bioanalyzer (Agilent, USA) was used to access the quality and size of the pre-processed and post processed samples and libraries.

### Routine genetic analysis

The FDA-approved Cobas mutation kit (Roche, Switzerland) was used to analyze the following mutations in the diagnostic biopsy tDNA: *KRAS* (G12S/R/C/V/A/D, G13D, Q61H, A146T), *NRAS* (Q61K/R/L/H), *BRAF* (V600E), and *PIK3CA* (E542K, E545K/G, Q546K, M1043I, H1047Y/R/L). The presence of the same mutations in the patient’s basal and on-treatment cfDNA samples was assessed by the highly sensitive BEAMing technique, as previously described^1^.

### Whole-exome sequencing

Sequencing libraries of cfDNA (15 ng), and gDNA and tDNA (70-110 ng) samples were prepared using the ThruPLEX Plasma-and DNAseq Kits (Rubicon Genomics Inc, USA), respectively. Barcode indices were added to samples during eight PCR cycles of template preparation, and 550 ng of each sample was processed through the SureSelectXT Target Enrichment System (Agilent SureSelect V5, ref. 5190-6208, protocol G7530-90000 version B1). xGen Blocking Oligos (IDT, Iowa, USA) were used as suggested by Rubicon Genomics. Captured targets were subsequently enriched by 11 cycles of PCR with KAPA HiFi HotStart (Kapa Biosystems), with a Tann of 60° and the following primers, which target generic ends of Illumina adapters: AATGATACGGCGACCACCGAGAT and CAAGCAGAAGACGGCATACGAGAT. For sequencing, magnetic bead-purified libraries with similar concentrations of cfDNAs and tDNA, and half the concentration of gDNA were pooled in order to increase coverage and favor the detection of non-inherited sub-clonal mutations. Sequencing was carried out in the Illumina HiSeq4000 platform. All sequencing data are going to be deposited in the European Nucleotide Archive (ENA) under the accession number ENA#202177, at the time of publication.

### Somatic mutation call

Bioinformatics analyses were performed using the NEXTGEN software (Softgenetics, USA), as previously described^2^. The detailed parameters used for the alignment and mutation call are provided as supplementary material. Briefly, FastaQ files were aligned using the BWA pipeline and the variants were processed by sequential stringent filters to exclude low-confidence variants. Only variants that passed the following filters were classified as high-quality and considered in the study: overall and allele scores ≥ 12; coverage ≥ 20; number of mutated reads ≥ 20; percentage of mutated reads ≥ 3% of cfDNA / tDNA and ≥ 35% of gDNA; F:R read balance ≥ 0.1; and F:R read percentage ≥ 0.45. The list of non-hereditary mutations detected by WES-cfDNA and WES-tumor was generated after disregarding germline variants (obtained by WES-gDNA). A detailed genomic annotation of the somatic mutations we identified, prediction of mutation pathogenicity based on predictor algorithms (SIFT, Polyphen2, LRT, Mutation Taster, Mutation Assessor, and other software packages included in the dbNSFP^3^), allele frequencies in population studies, such as 1000G and EXAC, and additional information are shown in Table S1.

### TaqMan SNP genotyping assay

A custom TaqMan^®^ genotyping assay for the detection of the *KDR* c.2518C (L840L) and *KDR* c.2518C>T (L840F) alleles was designed using the Thermofisher online Design Tool (oligonucleotides and probes are shown in Table S3).

### Genetic/protein database and protein structure analyses

Previously reported germline and somatic variants in *KDR* were retrieved from general population (EXACT^4^ and ESP^5^) and cancer (COSMIC^6^, GENIE^7^, PCAWGS^8^) sequencing public projects. The VEGFR2, EGFR, and ABL1 protein structures were obtained from the RCSB data bank; structurally analogous mutations in other cancer-relevant kinases were identified using MutationAligner^9^; kinase residues interacting with kinase inhibitors were mapped using the LigPlot^10^ software. Computational modeling of inhibitor binding to WT and L840F VEGFR2 was performed as previously described.

### Generation and treatment of the Avatar patient-derived xenograft (PDX) model

Liver metastasis biopsy was performed after tumor progression to capecitabine-bevacizumab rechallenge (Figures 1 and 1S). A fraction of the biopsy was used to generate the Avatar model as previously described by our group^11,12^. Expanded cohorts (five to six animals per arm) were treated with: anti-VEGF drugs (B20/murine and bevacizumab/human), VEGFR2 kinase inhibitors (axitinib, cabozantinib, cabozantinib:MEK inhibitor combo, lenvantinib, pazopanib, regorafenib, and sorafenib), and inhibitors of other kinases, such as afatinib (EGFR), crizotinib (MET), and MEK inhibitor (MAPK). Information on the treatment regimens is shown in Table S4.

### Cell lines

The human CRC cell lines used in the current study were selected based on their genotype in order to be as informative as possible for each experiment. Thus, we chose the Colo-320 cell line to interrogate the phenotypic changes caused by the overexpression of VEGFR2 mutants because it has the same genetic background as the patient´s tumor (mutated *TP53*/*APC* and WT *KRAS*/*BRAF*). The MDST8 CRC cell line was used for drug sensitivity studies because it naturally harbors the *KDR*/VEGFR2 R1032Q mutation, which we found to be a hot-spot VEGFR2 mutation in human cancers.

Colo320 and MDST8 colorectal cell lines were obtained from ATCC and cultured at 37°C in 5% CO_2_, in Roswelll Park Memorial Institute (RPMI) Medium 1640 + GlutaMAX (Gibco, USA) and Dulbeccos’ modified Eagle’s medium (DMEM) + 2 mM Glutamine (Gibco, USA), respectively, supplemented with 10% fetal bovine serum (FBS) (Thermo Fisher Scientific, USA). Porcine aortic endothelial (PAE) cell lines, kindly provided by Dr. Kurt Ballmer-Hofer, were grown in DMEM supplemented with 10% FBS.

### Generation of stable colorectal and endothelial cell lines

Colo-320 and PAE cell lines were used to generate cell lines stably expressing the VEGFR2 mutants using a previously described protocol^12^. Briefly, cells were seeded in 10-cm plates in the appropriate medium and were grown to 70% confluence. Transfection with constructs carrying either the empty vector or VEGFR2 (WT or mutant) was performed with polyethylenimine (PEI) as previously described^13^. Briefly, 30 μg of WT or mutant VEGFR2 plasmid (in the pBE vector containing the neomycine resistance gene, which confers resistance to the selection antibiotic G418) was mixed with 60 μl PEI (1 mg/ml in H_2_O) in 2 ml serum-free DMEM, incubated for 10 min at room temperature and added to the cells. Following a 3-h incubation at 37°C, the medium was changed, and the cells were allowed to grow to 100% confluence. Cells were re-seeded at a series of dilutions (1:1000-1:5000) in antibiotic selection medium (1 mg/ml G418) to allow for single colonies to grow, while non-transfected cells were dying. Individual colonies were consecutively transferred to 24-well and 6-well plates and screened by western blotting for VEGFR2 expression. To reduce polyclonality, colonies with the highest expression levels were subjected to 3 additional rounds of subcloning.

### VEGF stimulation and western blotting

Transiently transfected HEK293 cells or stable PAE cell lines expressing WT or L840F-KDR were starved in DMEM supplemented with 1% bovine serum albumin (BSA) for 4h at 37°C and were subsequently stimulated with 1.5 nM (60 ng/ml) VEGF_165_ for 10 min at 37°C. Following stimulation, the cells were scraped in lysis buffer (50 mM Tris pH = 8.0, 120 mM NaCl, 1% NP-40) supplemented with protease inhibitors (Roche, cat. Nr 04693159001) and phosphatase inhibitors (1 mM sodium orthovanadate and 20 uM phernylarsine oxide) and incubated for 30 min on ice. Cell lysates were collected as the supernanant of a centrifugation at 30,000 × *g* for 15 min and subjected to western blot analysis. The following antibodies were used to probe receptor activation: total KDR (Cell Signaling, cat. Nr 2479), phospho KDR at Y1175 (Cell Signaling, cat. Nr 2478). The secondary antibodies used were alkaline phosphatase (AP) conjugated (Southern Biotech). All antibodies were diluted at a 1:1000 ratio in 5% BSA in Tris-buffered saline, containing 0.05% Tween20 (TBST) buffer. The chemiluminescence signal was developed with the Novex AP Chemiluminescence substrate (Invitrogen, cat. Nr 100002906), recorded with an Amersham Imager 600 (Amersham), and quantified by ImageJ (NIH). Activation of KDR was assessed by the ratio of phospho-to-total signal.

### Tissue immunofluorescence

Immunofluorescence staining was performed to detect p-ERK and p-AKT. Formalin-fixed and paraffin-embedded tumors from Avatar models were cut into 3-mm-thick sections, deparaffinized, and preincubated with FBS to prevent nonspecific binding. The sections were incubated at room temperature for 30 min with a rabbit polyclonal antibody to p-ERK (1:300; Cell Signaling #9101) or a rabbit monoclonal antibody (D9E) to p-AKT (1:300, Cell Signaling #4060), followed by incubation with Alexa Fluor 555–conjugated donkey anti-rabbit IgG (1:400; Life Technologies#A27039) at 37°C for 20 min. Nuclei were counterstained with DAPI (Molecular Probes) at 1:1,000 dilution, and the slides were mounted with Mowiol 4-88 (Calbiochem). Images were acquired with a confocal TCS-SP5 (AOBS-UV) (Leica Microsystems) confocal microscope, equipped with a 20xHCX PL APO 0.7 N.A. objective.

### Proliferation assays

Proliferation assays were performed using the CellTiter-Glo^®^ Luminescent Cell Viability Assay (Promega). Briefly, cell lines were seeded in 96-well microtiter plates at a density of 10000 cells/well and were incubated for 24 h before adding the various drugs. A “mother plate” containing drugs at a concentration 200× higher than the final concentration to be used in the cell culture was prepared by serial dilutions of stock solutions of the drugs (10 mM) in DMSO. The appropriate volume from each drug (usually 2 μL) was added automatically (Beckman FX 96 tip) from this plate to the cell culture plate to reach the final concentration for each drug. Each concentration was assayed twice. The final concentration of DMSO in the tissue culture media did not exceed 1%. The cells were exposed to the drugs for 72 h and then analyzed using the CellTiter-Glo^®^ Luminescent Cell Viability Assay (Promega). Cell proliferation values were plotted against drug concentrations and fitted to a sigmoid dose-response curve using the Activity base software from IDBS in order to calculate growth inhibition (GI_50)_ values versus DMSO.

### Cloning and mutagenesis

The L840F and eight additional *KDR* mutations of interest identified on cancer databanks, as well as the K868M kinase-dead mutation, were generated by site-directed mutagenesis of WT *KDR*/VEGFR2 cloned on the pBE vector, using the QuikChange Kit (Agilent, USA) and the primers described in Table S5. Mutations were confirmed by Sanger sequencing of the entire open reading frame.

### Transfection and xenograft models

Colo-320 cell line-derived xenografts were generated from subcutaneous injections of 4 × 10^5^ cells resuspended in phosphate-buffered saline (PBS) in four nude mice per genotype. Tumors were measured weekly and the animals were sacrificed within two months or when tumors reached the established humane endpoint. Mice injected with empty vector or the K868M kinase-dead mutant were kept alive and monitored weekly for four months.

### Production of recombinant kinase domains of WT, L840F, and R1032Q VEGFR2

The kinase domains (residues 806-1171) of WT, L840F, and R1032Q VEGFR2 without the kinase insert domain (aa 940-989) were cloned, tagged with 6×His at their C-terminus, and expressed in the baculovirus-infected insect cell system. Proteins were purified by affinity chromatography on HisTrap columns, followed by size-exclusion chromatography on a HiLoad 16/600 Superdex 200 prep grade column (GE Healthcare, USA), using an ÄKTA system (GE Healthcare, USA). Fractions containing kinase domains were identified by sodium dodecyl sulfate polyacrylamide gel electrophoresis (SDS-PAGE) and concentrated by ultrafiltration up to 0.2 mg/ml. Protein mutations were confirmed by in-gel enzymatic digestion followed by liquid chromatography–mass spectrometry (LC-MS)/MS analysis.

### Biochemical Assays

The kinase activity of recombinant WT, L840F, and R1032Q VEGFR-2 (*KDR*) kinase domains, as well as that of a commercially available recombinant KDR cytoplasmic domain (residues 789-1356) (PV3660; ThermoFisher) that was used as a positive control were analyzed using the LANCE^®^Ultra time-resolved fluorescence resonance energy transfer (TR-FRET) assay from Perkin Elmer according to the manufactures’ instructions.. Briefly, the enzymes were titrated starting from an initial concentration of 5 µg/ml and proceeding with 1:4 serial dilutions, and were added to the reaction buffer (15 mM HEPES pH 7.4, 20 mM NaCl, 1 mM EGTA, 0.02% Tween 20, 10 mM MgCl_2_, 0.1 mg/ml BGG, 2 mM DTT), containing 15 μM ATP and 200 nM Ultralight^TM^-labeled Poly GT substrate in a total volume of 20 μl. The reaction was allowed to proceed in an Optiplate 384 from PerkinElmer for 60 min at room temperature. Reactions proceeded within the linear reaction time were then terminated by the addition of 20 mM EDTA and 4 nM Eu-W1024-labeled PY20 antibody. After an incubation of at least 60 min, the samples were excited with a Light Unit laser at 337 nm, and the emission of the LANCE Eu/APC (615/665 nm) was measured with an Envision reader (PerkinElmer). To test the effect of known VEGFR2 inhibitors on kinase activity, 0.3 ng WT VEGFR2 and 300 ng mutant VEGFR2 were used. The starting concentration of the inhibitors tested was 10 µM, followed by 1:5 serial dilutions. In order to calculate IC_50_ values of inhibition versus DMSO, the data were plotted against the inhibitor concentration and fitted to a sigmoid dose-response curve using the Activity base software from IDBS.

### Immunohistochemistry

Avatar tumor samples were fixed in 10% neutral buffered formalin (4% formaldehyde in solution) and paraffin-embedded. Subsequently, 3-μm-thick sections were cut from the samples, mounted in superfrost^®^plus slides, and dried overnight. Before staining, the sections were deparaffinized in xylene and re-hydrated through a series of decreasing ethanol concentration in water. Consecutive sections were stained with hematoxylin and eosin (H&E) and by immunohistochemistry, using an automated immunostaining platform (Ventana Discovery XT, Roche or Autostainer Plus Link 48). Antigen retrieval was first performed with high or low pH buffer (CC1m, Roche), endogenous peroxidase was blocked (3% hydrogen peroxide), and the slides were incubated with an anti p-ERK rabbit polyclonal primary antibody (1:300; Cell Signaling #9101) for 28 min. Subsequently, the slides were incubated with the corresponding visualization system (OmniRabbit, Ventana, Roche) with signal amplification conjugated with horseradish peroxidase. The signal was developed using 3,30-diaminobenzidine tetrahydrochloride (DAB) as a chromogen (Chromomap DAB, Ventana, Roche or DAB solution, Dako), while the nuclei were counterstained with Carazzi’s hematoxylin. Finally, the slides were dehydrated, cleared, and mounted with a permanent mounting medium for microscopic evaluation. The entire slide was scanned with a slide scanner (Axio Z1, Zeiss), and images were captured with the ZEN software (Zeiss) after evaluation by a trained veterinary pathologist. Image analysis and quantification were performed using the AxioVision software package (Zeiss).

### Kaplan-Meier analysis of mCRC patients

Kaplan-Meier survival data of 1,303 patients from four different cohorts of CRC patients with global gene expression and survival datasets available (GSE24551, GSE14333, GSE17538, GSE39582) were queried using the R2 microarray analysis and visualization platform (http://hgserver1.amc.nl/cgi-bin/r2/main.cgi). Concerning *KDR* gene expression in the combined datasets, 304 tumors had high levels of expression and 999 had low levels of expression.

